# Bubble Trouble: Conquering Microbubble Limitations in Contrast Enhanced Ultrasound Imaging by Nature-Inspired Ultrastable Echogenic Nanobubbles

**DOI:** 10.1101/633578

**Authors:** Al de Leon, Reshani Perera, Christopher Hernandez, Michaela Cooley, Olive Jung, Selva Jeganathan, Eric Abenojar, Grace Fishbein, Amin Jafari Sojahrood, Corey C. Emerson, Phoebe L. Stewart, Michael C. Kolios, Agata A. Exner

## Abstract

Advancement of ultrasound molecular imaging applications requires not only a reduction in size of the ultrasound contrast agents (UCAs) but also a significant improvement in the *in vivo* stability of the shell-stabilized gas bubble. The transition from first generation to second generation UCAs was marked by an advancement in stability as air was replaced by a hydrophobic gas, such as perfluoropropane and sulfur hexafluoride. Further improvement can be realized by focusing on how well the UCAs shell can retain the encapsulated gas under extreme mechanical deformations. Here we report the next generation of UCAs for which we engineered the shell structure to impart much better stability under repeated prolonged oscillation due to ultrasound, and large changes in shear and turbulence as it circulates within the body. By adapting an architecture with two layers of contrasting elastic properties similar to bacterial cell envelopes, our ultrastable nanobubbles (NBs) withstand continuous *in vitro* exposure to ultrasound with minimal signal decay and have a significant delay on the onset of *in vivo* signal decay in kidney, liver, and tumor. Development of ultrastable NBs can potentially expand the role of ultrasound in molecular imaging, theranostics, and drug delivery.

## Introduction

With an estimated 2.8 billion imaging procedures done in 2011 worldwide, ultrasound is among the most widely used and safest modalities for visualizing tissues and organs in real-time.^1^ *Therefore, a slight advancement on ultrasound technology can have a potentially large impact in the medical field*. As an alternative to commercially UCAs, nanodroplets, nanovesicles, and NBs have been studied. ^2–9^ Of growing interest is the use of NBs as UCAs for cancer detection, tumor characterization, image-guided surgeries and biopsies, as well as theranostics.^10–18^ Clinically-used UCAs (1 −10 μm microbubbles) are limited to intravascular applications due to their larger size.^12^ For both bubbles, one of the main limitations is the rapid *in vivo* signal decay; commercially available microbubble formulations have a signal half-life of under 2 min.^19–22^ Development of stable UCAs has been focused primarily on reducing surface tension, varying membrane components, counteracting Laplace pressure, preventing bubble coalescence, and utilizing hydrophobic gases.^15,23–27^ Still, most of the UCAs that show extended stability *in vitro* do not perform as well when continuously insonated or used *in vivo* because their shell is not designed to endure large deformations. Insonated UCAs undergo fast and extreme oscillations for which gas leaks out during expansion and shell materials shed off during compression.^28,29^ Furthermore, UCAs experience strong shear and turbulence as they circulate within the body.^30,31^ Ultrastable NBs can be designed by engineering the bubble membrane shell to be able to withstand deformations due to ultrasound and flow *in vivo*.

One particular membrane architecture that has been shown to redistribute stress, dissipate excess energy, and deform without irreversible damage is a bilayer with contrasting elastic properties. Membranes comprised of contrasting elastic properties have been shown to provide synergistic improvement in toughness and resilience against defects, stress, and deformations.^32–34^ In nature, this architecture can be found not only in the bacterial cell envelope, but also apparent in the design of interfaces in mammalian blood vessels, skin, and red blood cells.^35,36^ We hypothesize that adapting this architecture can impart NBs an enhanced capacity to minimize gas leakage and membrane shedding during insonation and therefore provide better persistence as it circulates within the body.

Development of ultrastable NBs can facilitate diagnostic ultrasound imaging of various malignancies by providing more accurate, less transient visualization of tumor vasculature and with appropriate molecular targets, detection of cellular biomarkers. In addition, for molecular imaging, stable echogenic NBs can enable a more reliable extravasation beyond the tumor vasculature. Likewise, for drug delivery, a more robust shell can prevent pre-mature release of loaded drug far from the target sites brought about by the collapse of NBs in circulation. Here, we engineered the membrane of ultrastable NBs to be composed of a compliant phospholipid layer by incorporating propylene glycol (PG), an edge-activator used in ultra-deformable liposomal formulations, and glycerol (Gly), a membrane stiffener known to increase buckling of lipid monolayers (Figure 1a).^37–41^ The unique interfacial design reduces *in vitro* signal decay of NBs under continuous exposure to ultrasound of varying strength and pulse repetition frequencies, enabling a substantial improvement in both *in vivo* half-life and *in vivo* signal decay onset as compared to a common commercially available UCAs.

**Figure 1.**
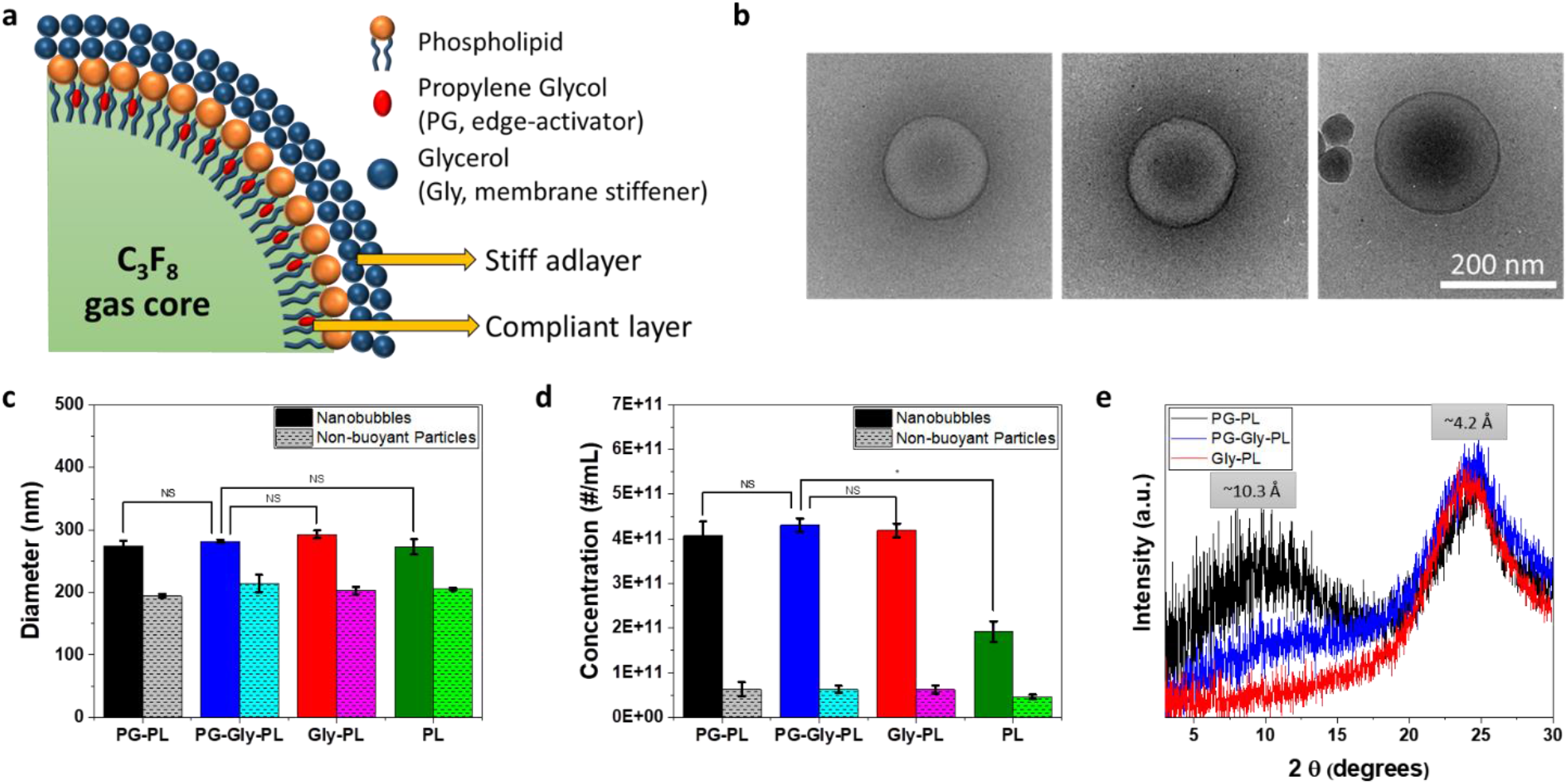
Schematic, size, concentration, and structural property of NBs. **(a)** Schematic representation of the membrane of the ultrastable NB showing a bilayer architecture with elastic contrast. **(b)**. Cryo-electron microscopy (cryo-EM) images of PG-Gly-PL showing the nanobubble membrane and the dense C_3_F_8_ gas core. Diameter **(c)** and concentration **(d)** measurement of NB via a resonant mass measurement capable of differentiating buoyant (NBs) and non-buoyant particles (micelles, liposomes, etc.). Mean ± SE (n = 5) **(e)** X-ray diffractogram of the membrane of the freeze-dried NBs. The NBs were isolated and freeze-dried overnight. Peak corresponding to d = 4.5 Å is observed for all NB but only PG-PL and PG-Gly-PL have a peak corresponding to d = 10.3 Å implying the incorporation of PG into the phospholipid membrane.

## Results and Discussion

NBs containing a perfluoropropane (C_3_F_8_) gas core were prepared by dissolution of a phospholipid mixture in PBS solution containing PG and Gly (**PG-Gly-PL**), followed by activation via amalgamation and isolation by differential centrifugation.^26,27^ To directly compare the effects of the glycerol and propylene glycol we also prepared phospholipid stabilized NBs without glycerol (propylene glycol only (**PG-PL**)), without propylene glycol (glycerol only (**Gly-PL**)), or using phospholipids alone (**PL**) without additives. Cryo-EM images shown in Fig 1b and S2 show a dense C_3_F_8_ gas core enclosed by a phospholipid membrane.^42^ Particle size distribution and concentration for each NB type were determined through resonant mass measurement capable of distinguishing buoyant and non-buoyant particles (liposomes, micelles, etc).^43–47^ The average diameter of NB was measured to be 274 ± 8 nm, at a concentration of 4.07 x 10^11^ ± 3.15 x 10^10^ NBs/mL, with no particles larger than 1 μm observed in the population (Figure 1c, 1d, S1). Cryo-electron microscopy confirms an average diameter of ^~^200 nm and reveals the nanobubble membrane and dense C_3_F_8_ gas core (Figures 1b and S2). Aside from NBs, non-buoyant particles were also measured to be 194 ± 3 nm in average diameter and concentration of 6.32 x 10^10^ ± 1.58 x 10^10^ particles/mL. These non-buoyant particles, which are invisible under ultrasound, were reported to sterically hinder coalescence of UCAs, thereby contributing to their overall stability.^48^ NBs in PBS showed a slightly negative charge as measured via zeta potential, with no significant differences among formulations (Table S2). Although the isolated NBs have similar size, charge, and concentration, analysis of the shell material by X-ray diffraction (XRD) showed an additional broad amorphous peak centered at 2θ = ^~^10° for **PG-PL** and **PG-Gly-PL** (Figure 1e) suggesting that PG expands the distance between phospholipids from ^~^4.2 Å to ^~^10.3 Å at some locations.^49,50^ This is consistent with the reported incorporation of edge-activators such as PG into membranes of ultradeformable liposomes. These ultradeformable liposomes have been reported to deform as they pass through a narrow channel (*e.g*. skin pore) and reform without disruption of their vesicular structure.^37,38,51–54^ Of note, the broad peak centered at 2θ = ^~^10° is not present for pure components (Figure S3a) and no preferred crystalline orientation is observed in 2D XRD (Figure S3b-d).^55^

The ability of a NB to function as an UCA depends on the acoustic impedance difference between the gas and its surrounding and the nonlinearity of the NB oscillation.^56–58^ Tissues do not exhibit nonlinear scattering of the ultrasound wave, enabling high contrast, which is ideally suited for molecular imaging. Therefore, understanding the nonlinear behavior of NB oscillation can facilitate better parameter optimization for molecular imaging and theranostics.^59^ Subharmonic, ultraharmonic, second harmonic, as well as fundamental harmonic oscillations, were observed when single NBs were insonated by a 100% bandwidth 25 MHz ultrasound transducer and pressure amplitude of 300 kPa as shown in Figure 2a. Dilute suspensions of NBs were introduced in PBS, and the scattering of single NBs were recorded following a procedure described elsewhere.^60,61^ **PG-PL**, with the most flexible shell, shows the largest oscillation amplitude for both fundamental (f) and second harmonic (2f) oscillations. This is consistent with the Marmottant model, which predicts that the backscatter intensity is directly proportional to shell flexibility.^62–64^ In contrast, **Gly-PL** shows the lowest oscillation amplitude for fundamental and second harmonic, suggesting that it has a stiffer shell than **PG-PL**. The interaction between Gly and phospholipids has been well-studied experimentally, numerically, and via molecular dynamics. ^39,40,65–68^ Recently, Terakosolphan, et al. found via molecular dynamics simulation that glycerol preferentially interacts with the phosphate and choline in the headgroup of phospholipids. Furthermore, these are experimentally confirmed via Langmuir pressure-area isotherms, FT-IR spectroscopy, fluorescence-based membrane transition temperature determination, and small angle neutron scattering.^39^ Pocivavsek et al., in particular, reported the formation of a 10 Å-thick Gly-enriched vitrified adlayer at the phospholipid/solvent interface as measured by neutron and X-ray reflectivity.^41^ In the same study, it was shown that the Gly adlayer stiffens the phospholipid monolayer at the air-water interface enabling its folding/buckling normal to the interface when compressed. This stiffening, caused by inter- and intra-hydrogen bonding with the phospholipid, limits the oscillation of **Gly-PL** NBs when exposed to ultrasound field. **PG-Gly-PL**, which has a membrane stiffness between that of **PG-PL** and **Gly-PL**, shows intermediate amplitude for both fundamental and second harmonic. Furthermore, the **PG-Gly-PL** NBs undergo a highly stable oscillation with minimal change in amplitude with time. Aside from fundamental and second harmonic, subharmonic (f/2) and ultraharmonic (3f/2) signals can also be detected for all samples further confirming the non-linear nature of oscillation of NB.

**Figure 2.**
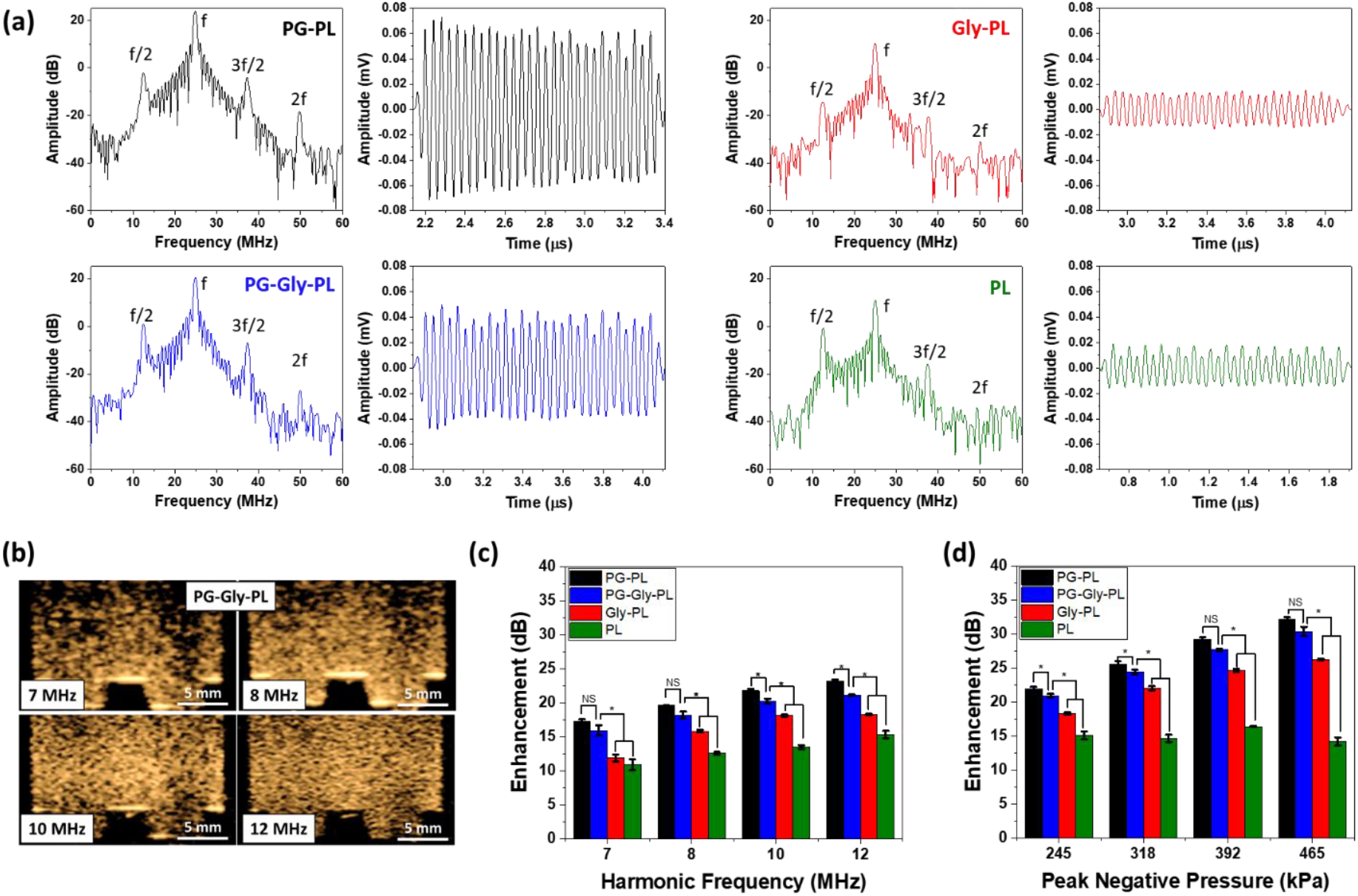
Nonlinear oscillation of a single NB and a solution of NBs. **(a)** Representative nonlinear backscatter from a single NB from a narrowband 30 cycle ultrasound pulse and the corresponding frequency spectrum (25 MHz transducer at 300kPa). Presence of subharmonic (f/2), ultraharmonic (3f/2), and second harmonic (2f), in addition to its fundamental (f) harmonic signal in the frequency spectra confirms the nonlinear oscillation of NBs. Minor changes in backscattered amplitude during the 30 cycle pulse shows how stably **PG-Gly-PL** NB oscillates when insonated by 25 MHz ultrasound. The magnitude of amplitude is consistent with the predicted stiffness of the NB shell having **PG-PL** as the most flexible, **Gly-PL** as the stiffest, and **PG-Gly-PL** with intermediate stiffness. **(b)** Contrast harmonic images (2^nd^ harmonic) of **PG-Gly-PL** at different receiving frequency showing improvement in both intensity and resolution of the backscattered signal. NB with concentration of 4 x 10^9^ NBs/mL was placed in a tissue-mimicking agarose phantom. Only the second harmonic backscattered signal is visible in contrast harmonic imaging mode which implies that the backscattered signal from the agarose phantom, similar to tissues, does not have non-linear component. Enhancement of nonlinear backscattered signal of NB (concentration of 4 x 10^9^ NB/mL) relative to tissue-mimicking agarose phantom at different receiving frequency **(c)** and peak negative pressure **(d)**.

*In vitro* echogenicity of a solution of NBs (concentration of 4 x 10^9^ NB/mL) relative to a tissue-mimicking agarose phantom (Figure S4a) was studied by analyzing the nonlinear backscatter at different ultrasound frequencies using a clinical scanner in contrast harmonic imaging mode (Figure S5). The enhancement, as well as the resolution, increased as the receiving frequency is changed from 7 MHz to 12 MHz (Figure 2b and 2c). As expected, the agarose phantom did not show any nonlinear acoustic activity, similar to tissues and organs, as shown by dark areas in Figure 2b.^69^ Consistent with what has been observed when single NB were insonated (Figure 2a), the flexible **PG-PL** bubbles showed the greatest enhancement while, the stiff **Gly-PL** showed a lower enhancement suggesting lower magnitude of oscillation as compared to **PG-PL**. **PG-Gly-PL** has intermediate enhancement because it has both a compliant and a stiff adlayer and thus intermediate enhancement. A similar trend is observed as the peak negative pressure of the transmitted ultrasound (Figure 2d and S6) is increased with **PG-PL** showing the greatest enhancement.

As stated above, one of the main challenges with current UCAs is rapid signal decay due to dissipation of the gas from the bubble core, which is immensely amplified when the UCAs are continuously oscillating due to ultrasound. To assess bubble *in vitro* stability under constant insonation, we placed the NB solution in a tissue-mimicking phantom with a narrow channel (Figure S4b) and exposed it to ultrasound at varying frame rates and pressures (Figure S7 and S8). A narrow channel was specifically used to ensure that the NBs were continuously insonated, in contrast to the typical stability test wherein the NB can move in and out of the ultrasound field (Figure S4b).^22,70^ In these experiments, **PG-Gly-PL** showed the slowest signal decay (Figure 3a, b) and the longest *in vitro* half-life for all pressures and frame rates (Figure 3c, d). The synergistic effect of having a bilayer with elastic contrast designed similar to natural membranes enabled the **PG-Gly-PL** to resist failure under continuous oscillation. We hypothesize that while PG introduces elastic and compliant defects in the shell that absorbs excess energy during compression as demonstrated with XRD in Figure 1d, Gly forms a vitrified adlayer that provides stiffness, limits strain during expansion, and functions as an additional layer resisting gas diffusion.^41,71,72^ Furthermore, **PG-PL** has a longer *in vitro* half-life than **Gly-PL** suggesting that having a compliant shell imparts better stability under ultrasound than having a stiff shell. For all cases, **PL** always showed the lowest enhancement and shortest *in vitro* half-life, likely brought about by its inability to endure oscillating deformations. Interestingly, and similar to prior observations by our group, the equilibrium surface tension appeared to have a lesser influence on the bubble stability in this system as **Gly-PL** or **PG-PL** showed the lowest surface tension (but *not* the highest stability) when measured via pendant drop or rising bubble with perfluoropropane gas, respectively (Figure S9).^26^

**Figure 3.**
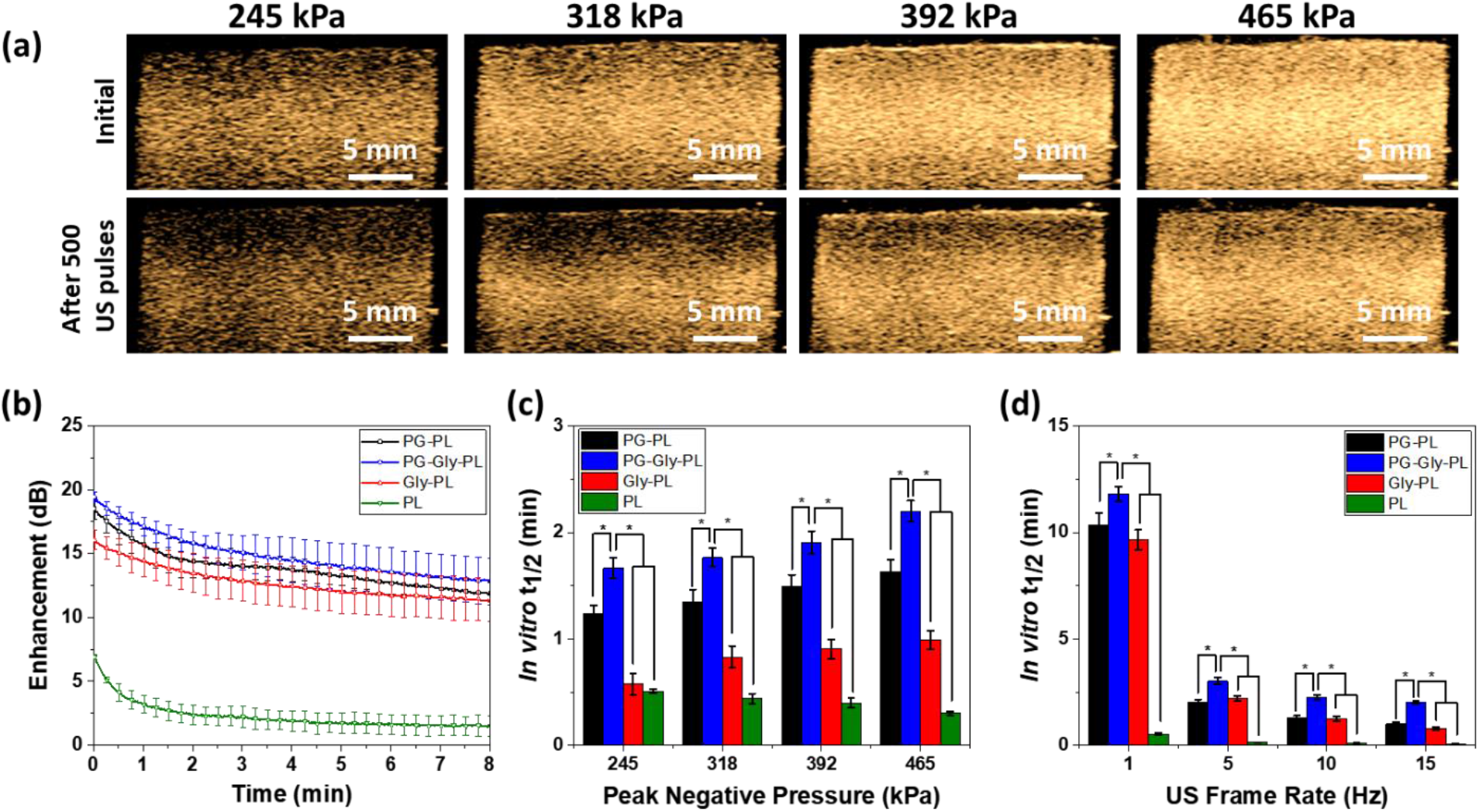
In vitro characterization of NBs in tissue mimicking agarose phantom. **(a)** Contrast harmonic images of **PG-Gly-PL** (concentration of 4 x 10^9^ NB/mL) before and after application of 500 frames of ultrasound at different peak negative pressure (P = 245 kPa to 465 kPa) at 1 frame per second. A dilute solution of **PG-Gly-PL** NB is placed in a tissue-mimicking agarose phantom with a thin channel (1 mm) to ensure minimal diffusion of NB in and out of the ultrasound field. **(b)** Representative signal decay curve of NBs continuously exposed to 245 kPa pressure at 1 frame per second. Mean ± SE (n = 3). **(c)** *In vitro* half-life (t_½_) of nonlinear backscattered signal dilute NBs placed in a tissue mimicking agarose phantom and exposed to ultrasound with varying peak negative pressure at 15 frames per second. Mean ± SE (n = 3) **(d)** *In vitro* half-life of nonlinear backscattered signal of dilute NBs placed in a tissue mimicking agarose phantom and at 245 kPa pressure and varying frame rate. Mean ± SE (n = 3).

Development of long-circulating NBs can have a significant impact in advancing the role of ultrasound in applications such as molecular imaging, and can open the door to more effective targeted theranostic or drug-delivery applications in cancer and a number of other diseases where nanomedicine already plays a critical role.^73,74^ Clinical practice with current UCAs is limited to circulation times of ^~^5 mins, which is the reported *in vivo* lifetime of Definity^®^ and Sonovue^®^, as UCAs quickly dissipate and are consumed by macrophages.^22^ The *in vivo* lifetime of current UCAs, even with a smaller, sub-micron footprint, is thus insufficient to enable reliable extravasation and retention in tumors. To confirm if the observed *in vitro* stability of **PG-Gly-PL** NBs is translatable to *in vivo*, we injected our NB solution into the tail vein of a mouse and continuously acquired images of the kidney and liver for 30 mins with a clinical ultrasound in contrast harmonic imaging mode (Figure 4a, b). For comparison, we performed an identical experiment with a commercially-available and FDA-approved UCA (**Lumason**^®^). All samples show improvement in kidney contrast within 1 min after injection, but the signal from **Lumason**^®^ dissipated within 5 min with *in vivo* half-life (t_1/2_) of 1.56 ± 0.27 min. The *in vivo* t_1/2_ for **Gly-PL** was determined to be 14.24 ± 1.08 min, **PG-PL** to be 12.19 ± 1.2 min, and **PG-Gly-PL** to be 18.32 ± 0.40 min. Despite having similar maximum non-linear image enhancement, **PG-Gly-PL** showed a 12x longer *in vivo* half-life than **Lumason**^®^ (Figure 4c and d). The enhancement of **PG-Gly-PL** was similar to **PG-PL** and higher than **Gly-PL**, but **PG-Gly-PL** bubbles showed a longer *in vivo* half-life than both **PG-PL** and **Gly-PL**. This indicates that **PG-Gly-PL** is not only the most stable under continuous ultrasound exposure *in vitro*, but also at higher temperature, under shear and turbulence in physiological blood flow, and in the presence of macrophages *in vivo*. In is interesting to note that the circulation time of bubbles without glycerol was shorter than both bubbles formulated with glycerol. This could also be indicative of reduced RES clearance due to the presence of glycerol on the bubble surface, an effect previously explored using liposomes.^75^ Furthermore, while **Lumason**^®^ followed the two-compartment pharmacokinetic model typical for UCAs (Figure 4b), the NBs followed a logistic behavior wherein the intensity remained at its maximum for some time before it started to decay (*in vivo* decay onset).^76^ The *in vivo* decay onset (Figure 4e) of **PG-Gly-PL** was significantly longer than **Lumason**^®^ (20x longer), **PG-PL**, and **Gly-PL**. This implies that continuous acquisition of high quality images can be performed in preclinical animal models for over 15 min following only a single bolus injection, which is not possible with **Lumason**^®^ as its signal immediately decay after reaching the peak. Similar behavior was also observed in the liver despite the presence of stellate macrophages known to efficiently eliminate foreign objects (Figure S10).

**Figure 4.**
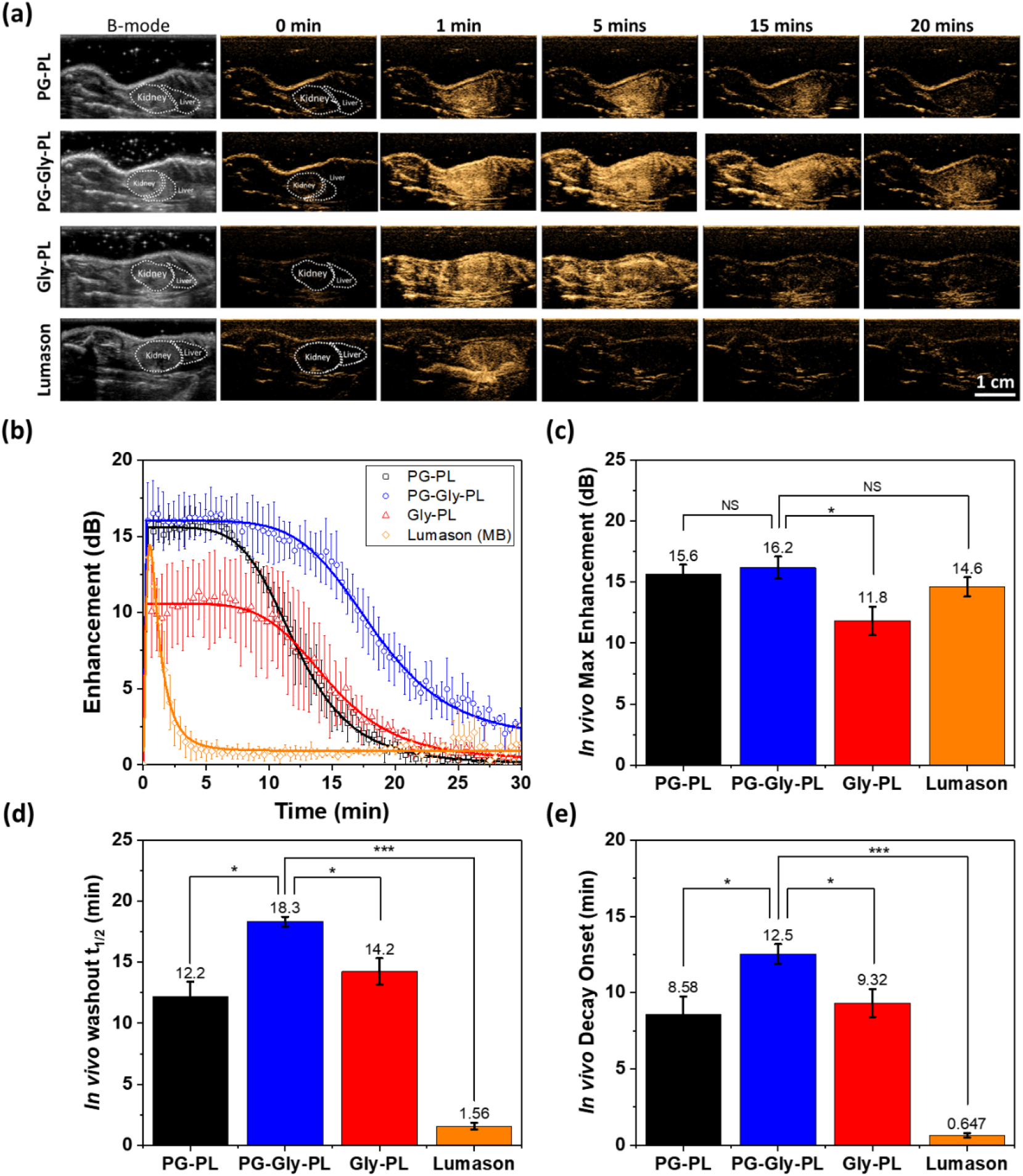
*In vivo* characterization of NBs in mouse kidney stability relative to commercially available UCA in mice. **(a)** NBs and commercially available UCA (Lumason) were injected via tail-vein and a cross-section of the kidney and liver was imaged at 12 MHz, 245 kPa pressure, and 0.2 frames per second. Left: Representative B-mode images of the kidney and liver before the injection of UCAs. 0 min-20 mins: Series of images showing the signal onset and signal decay of various UCAs at different time points. Minimal to no contrast can be observed before injection (t= 0) which is expected because backscatter signals from kidney and liver do not have any nonlinear properties. **(b)** Representative signal decay curve of NBs and Lumason^®^ showing the delayed signal decay onset and longer *in vivo* half-life of PG-Gly-PL. **(c)** *In vivo* maximum enhancement, **(d)** *in vivo* washout half-life, and **(e)** *in vivo* decay onset of NBs and Lumason extracted from the enhancement vs time curve.

To further demonstrate the unique capabilities of the **PG-Gly-PL** for practical applications, we performed a similar imaging protocol in a mouse with flank colorectal tumor (Figure 5). For comparison, a parallel experiment was performed with **Lumason**^®^. In addition, as an internal control, we also monitored the contrast dynamics in the kidney for both agents. Consistent with data shown in Figure 4, **Lumason**^®^ enhances both tumor and kidney a minute after injection, but rapidly dissipates within 5 minutes (Figure 5a-c). In contrast, **PG-Gly-PL** enhances the contrast of the kidney immediately, with *in vivo* decay onset at 19.67 ± 2.56 min and *in vivo* half-life of 24.38 ± 3.39 min (Figure S11). It is also worth noting that although the *in vivo* max enhancement of **Lumason**^®^ (16.94 ± 0.27 dB) in kidney is not significantly different than **PG-Gly-PL** (16.17 ± 0.86 dB), **PG-Gly-PL** (11.59 ± 1.11 dB) shows a 2-fold increase in *in vivo* max enhancement in tumor as compared to **Lumason**^®^ (5.14 ± 1.10 dB) (Figure S12).

**Figure 5.**
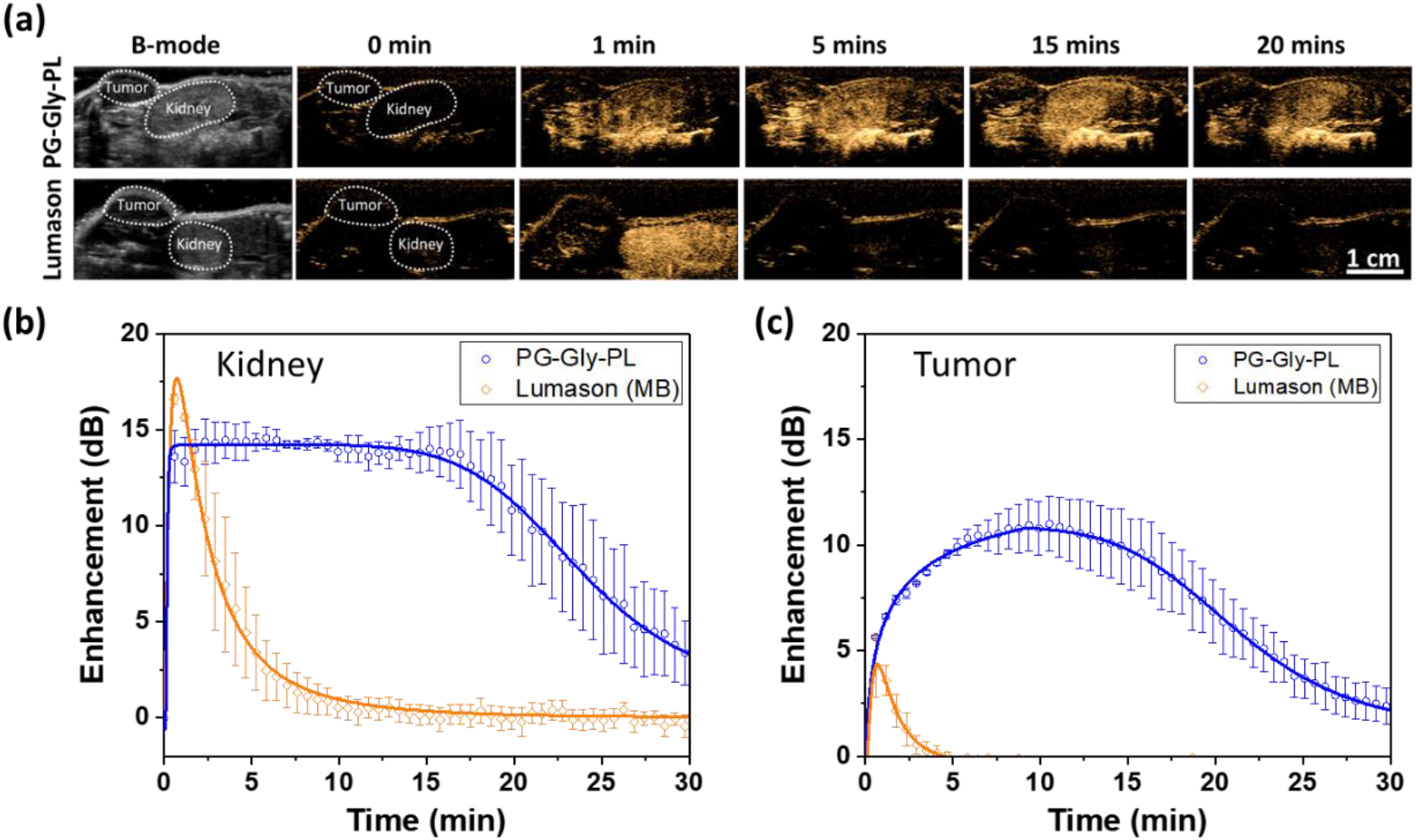
*In vivo* characterization of NBs in mouse kidney and flank colorectal tumor stability relative to commercially available UCA (Lumason^®^) in mice. **(a)** NBs or Lumason^®^ were injected via tail-vein and a cross-section of the kidney and flank tumor was imaged at 12 MHz, 245 kPa pressure, and 0.2 frames per second. Left: Representative B-mode images of the kidney and tumor before the injection of UCAs. 0 min-20 mins: Series of images showing the signal onset and signal decay of various UCAs at different time points. Minimal to no contrast can be observed before injection (t= 0) which is expected because backscatter signals from kidney and tumor do not have any nonlinear properties. Representative signal decay curves of **PG-Gly-PL** and Lumason^®^ in kidney **(b)** and in tumor **(c)** showing the delayed signal decay onset and longer *in vivo* half-life of **PG-Gly-PL**.

**Figure 6.**
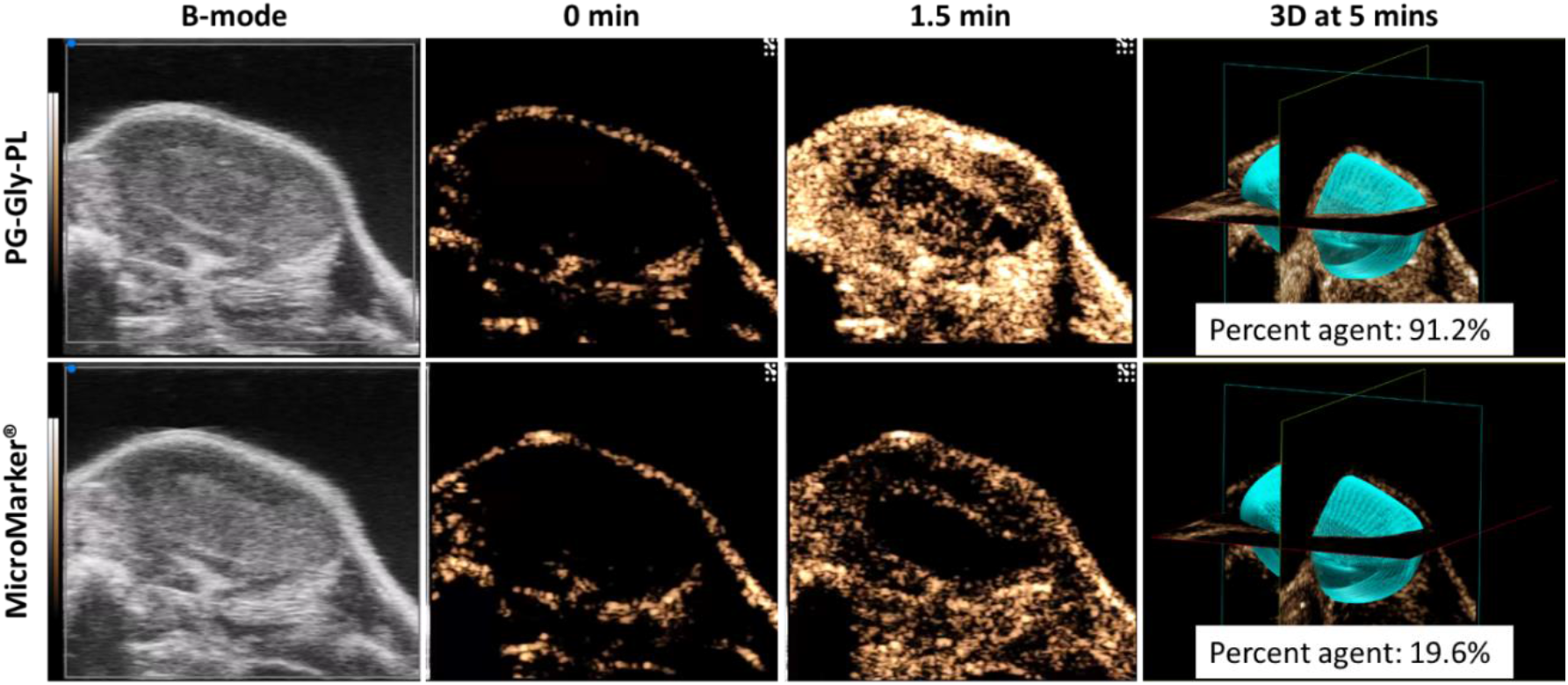
*In vivo* characterization of **PG-Gly-PL** in flank colorectal tumor stability relative to **MicroMarker**^®^ in mice showing comparison on the extent of UCA filling of tumor both in 2D and 3D. Tumor volume: 229.975 mm^3^.

Although the use of clinical ultrasound equipment demonstrates the potential for clinical translation of ultrastable NB, the equipment is not ideal for imaging small mouse tumors. We therefore examined performance of **PG-Gly-PL** compared to **MicroMarker**^®^ research microbubble (FujiFilm VisualSonics) optimized for use with a research ultrasound scanner (VisualSonics Vevo 3100). For this experiment, we injected **PG-Gly-PL** or **Micromarker**^®^ into a mouse with colorectal tumor and captured 2D contrast images for 5 mins before performing a 3D scan of the tumor. For both experiments, the transducer positon was held constant for a more direct comparison. Injection of **PG-Gly-PL** led to significantly higher enhancement at 1.5 min that highlights the entire tumor volume (outer shell and inner core), in comparison to **MicroMarker**^®^ that highlights mainly the periphery of the tumor. Videos of the contrast dynamics of both **PG-Gly-PL** and **MicroMarker**^®^ can be seen in supplementary information as videos 1 and 2, respectively. 3D volume analysis of the presence of contrast signal in the entire tumor volume shows >4-fold increase in coverage with **PG-Gly-PL** nanobubbles. Expanding the use of ultrasound in these applications could make crucial diagnostic imaging tools significantly more accessible to a broader global population as compared to modalities such as magnetic resonance imaging (MRI) and computed tomography (CT). Colorectal cancer, for example, is the second leading cause of cancer-related deaths for both men and women in the United States,^77,78^ yet the use of ultrasound for colorectal cancer diagnosis has been essentially abandoned by current guidelines in favor of computed tomography (CT).^79^ However, there has been recent interest in assessing colorectal tumor angiogenesis by quantitative contrast-enhanced endoscopic ultrasound.^80^ New contrast agents could provide complementary data in this area and eventually lead to a more effective diagnosis.

## Conclusions

We have shown that our ultrastable NB (**PG-Gly-PL**) with membrane structure similar to bacterial cell envelope has minimal signal decay when insonated continuously *in vitro*, has longer *in vivo* half-life, and has a delayed onset of *in vivo* signal decay. Despite having a bilayer structure with contrasting elastic property, the backscattered ultrasound from the ultrastable NB still has nonlinear properties when insonated as a single NB or as population. The enhanced *in vivo* stability of the ultrastable NB can advance the use of ultrasound in molecular imaging, theranostics, and drug delivery. Compared to **Lumason**^®^ (FDA-approved UCA) and **MicroMarker**^®^, our ultrastable nanobubble is able to more effectively enhance the entire tumor vasculature for an extended duration, making it the ideal potential agent for enabling future molecular imaging applications of ultrasound.

## Experimental Section

### Preparation and Isolation of NB solution

**PG-GLy-PL** solution (10 mg/mL) was prepared by first dissolving 1,2-dibehenoyl-sn-glycero-3-phosphocholine (C22, Avanti Polar Lipids Inc., Pelham, AL), 1,2 Dipalmitoyl-sn-Glycero-3-Phosphate (DPPA, Corden Pharma, Switzerland), 1,2-dipalmitoyl-sn-glycero-3-phosphoethanolamine (DPPE, Corden Pharma, Switzerland), and 1,2-distearoyl-snglycero-3-phosphoethanolamine-N-[methoxy(polyethylene glycol)-2000] (ammonium salt) (DSPE-mPEG 2000, Laysan Lipids, Arab, AL) into propylene glycol (PG, Sigma Aldrich, Milwaukee, WI) by heating and sonicating at 80 °C until all the lipids were dissolved. Mixture of glycerol (Gly, Acros Organics) and phosphate buffer solution (0.8 mL, Gibco, pH 7.4) preheated to 80 °C was added to the lipid solution. The resulting solution was sonicated for 10 mins at room temperature. Note that we typically prepared batches of 5 or 10 samples at a time and the amounts were adjusted as such. The solution (1 mL) was transferred to a 3 mL headspace vial, capped with a rubber septum and aluminum seal, and sealed with a vial crimper. Air was manually removed with a 30 mL syringe and was replaced by injecting octafluoropropane (C_3_F_8_, Electronic Fluorocarbons, LLC, PA) gas. **PG-PL** and **Gly-PL** solutions were prepared similar to **PG-Gly-PL** but with 1 mL of PG and 1 mL of Gly, respectively, instead of 0.5 mL of PG and 0.5 mL of Gly. **PL** solution was prepared by directly dissolving the phospholipid mixture into 1 mL PBS. After air was replaced by C_3_F_8_, the phospholipid solution was activated by mechanical shaking with a VialMix shaker (Bristol-Myers Squibb Medical Imaging Inc., N. Billerica, MA) for 45s. Nanobubbles were isolated from the mixture of foam and microbubbles by centrifugation at 50 rcf for 5 mins with the headspace vial inverted, and the 100 μL NB solution withdrawn from a fixed distance of 5 mm from the bottom with a 21G needle.

### Cryo-EM of NBs

Samples of the PG-Gly-PL NBs washed with PBS after isolation at concentrations around 10^10^ particles were applied to Quantifoil holey carbon EM grids (R2/2, 400 mesh:EMS). The EM grids were glow discharged for 30 sec at 15 mA before the sample was applied. 4 μL aliquots of the NB solution were applied to the EM grids and blotted for 6 sec. Following application, the grids were plunge frozen using a Vitrobot. The prepared cryoEM grids were imaged using a ThemoFisher/FEI TecnaiSpirit cryo-transmission electron microscope (operated at 80kV) equipped with a Gatan US4000 CCD camera. Micrographs were collected at 30kX magnification with a defocus range of −2.5 to −3 μm.

### Size, concentration, and surface charge of NBs and non-buoyant particles

The size distribution and concentration of NBs were characterized with a Resonant Mass Measurement (Archimedes^®^, Malvern Panalytical) equipped with a nanosensor capable of measuring particle size between 50 nm and 2000 nm. The nanosensor was pre-calibrated with 565 nm polystyrene beads (Thermo Scientific™ Nanospehere Size Standards 3560A). The NB solution was diluted with PBS (500x) to obtain an acceptable limit of detection (< 0.01 Hz) and coincidence (< 5%). Prior to any measurement, a 5-min PBS blank was run to ensure that the system fluidics and sensor were free of particles. Also, between measurements the sensor and microfluidic tubing were rinsed for 30 seconds with PBS followed by 2 “sneezes” for at least 3 cycles. During the sample measurement, NB solution was loaded for 120 seconds and analyzed at 2 and 5 psi, respectively. Samples measurement was finalized after 1000 particles were measured. Data was exported from the Archimedes software (version 1.2) and analyzed for positive and negative counts. Dilution was accounted for in calculating the NB concentration. Surface charge of the diluted NB solution (500X) was measure with an Anton Paar Litesizer™ 500.

### Crystallinity analysis of NB membrane by x-ray diffractometer (XRD)

1D and 2D X-ray diffraction experiments were performed using a Bruker Discover D8 X-ray diffractometer, with a monochromated X-ray source (used with a Co Kα X-ray tube), configured in point focus mode. Isolated NB solution was transferred to a 1.5 mL Eppendorf microcentrifuge tube and was flashed frozen with liquid nitrogen before being loaded to a freeze-dryer (Labconco Freezone 2.5) set at – 50 °C. The NB solution was freeze-dried for at least 48 hours before placing on a zero diffraction plate sample holder for XRD.

### Nonlinear oscillation of a single NB

The backscatter signal from single NB events were captured through insonation of a dilute solution of NBs, with continuous pulse trains of 30 cycles at 25 MHz frequency for pressures value of 300 kPa. The ultrasonic signals were transmitted and received by a Vevo 770 ultrasound instrument (VisualSonics Inc. Toronto, Ontario). The transducer has a 20 MHz center frequency and 100% bandwidth. The dilute solution enabled the recording of single scattering events from individual NBs within the focus of the ultrasound transducer, as has been shown previously with cells and microspheres

### Echogenicity of NB solution

Nonlinear ultrasound imaging was done on AplioXG SSA-790A clinical ultrasound imaging system (Toshiba Medical Imaging Systems, Otawara-Shi, Japan) with a 12 MHz center frequency linear array transducer (PLT-1204BT). Images were acquired in contrast harmonic imaging (CHI) mode with parameters set as: 65 dB dynamic range, 70 dB gain, receiving frequency (7, 8, 10, and 12 MHz) and peak negative pressure (245, 318, 382, 465 kPa) depending on the experiment. NB solution was diluted (1mL, 100x diluted, concentration of 4 x 10^9^ NBs/mL) to avoid signal attenuation and was placed in a tissue-mimicking agarose phantom. The agarose phantom was composed of 1.5 wt% agarose in Milli-Q water (resistivity of ^~^18 MΩ·cm) heated in a microwave until the agarose is dissolved. The hot agarose solution was then poured into a mold avoiding any trapped bubbles and cooled down to obtain phantom with the desired channel dimension (Figure S4a). Intensity of the backscattered nonlinear ultrasound signal was determined using a pre-loaded quantification software (CHI-Q). Enhancement by NBs was calculated by normalizing the measured backscattered ultrasound intensity of the NB solution with respect to the backscattered ultrasound intensity of the tissue-mimicking agarose phantom.

### Stability of NB under continuous insonation

NB solution was diluted (400 μL, 100x, concentration of 4 x 10^9^ NBs/mL) with PBS and was placed in a tissue-mimicking agarose phantom with a thin channel (L x W x H = 22 x 1 x 10 mm) as shown in Figure S4b. Thin channel was chosen to ensure that the diffusion of NB in and out of the ultrasound field is minimized and NBs were continuously insonated. Nonlinear ultrasound imaging was done on AplioXG SSA-790A clinical ultrasound imaging system (Toshiba Medical Imaging Systems, Otawara-Shi, Japan) with a 12 MHz center frequency linear array transducer (PLT-1204BT). Images were acquired in contrast harmonic imaging (CHI) mode with parameters set as: 65 dB dynamic range, 70 dB gain, imaging frame rate (1, 5, 10, 15 fps) and peak negative pressure (245, 318, 382, 465 kPa) depending on the experiment. 500 frames of ultrasound images were acquired and the intensity per frame is analyzed with a built-in software (CHI-Q). Enhancement by NBs was calculated by normalizing the measured backscattered ultrasound intensity of the NB solution with respect to the backscattered ultrasound intensity of the tissue-mimicking agarose phantom. In vitro half-life (in vitro t_1/2_) was calculated by using the calculated enhancement and with the formula for first order decay.

### Surface tension measurement of NB solution

Surface tension via pendant drop method was performed by dispensing a drop of the lipid solution at the tip of a microsyringe (SY20, Kruss Scientific, Germany) with a 0.5 mm outer diameter needle (Krüss, model: NE31). The needle used is made with stainless steel with PTFE coating and with stainless steel luer-lock connector. Images of the drops (n=3) was acquired with a KSV CAM 200 Optical Contact Angle Meter. Surface tension was calculated using the built-in software by analyzing the dimension of the drop and fitting into the Young-Laplace equation. Surface tension via rising drop method with air is done by dispensing an air bubble at the tip of a J-shaped needle (Krüss, model: NE71, 0.484 mm diameter) immersed in a lipid solution. Images of the bubbles (n = 3) was acquired and analyzed similar to the previous technique. Surface tension using rising bubble method with C_3_F_8_ was done similarly but replacing air with C_3_F_8_.

### NB and Lumason^®^ stability *in vivo* studies

Animal experiments were conducted in compliance with the Institutional Animal Care and Use Committee (IACUC) at Case Western Reserve University. In all procedures, the animals were anesthetized with 3% isoflurane with 1 L/min oxygen. Both NB solutions and Lumason (sulfur hexafluoride lipid-type A microspheres, Bracco Diagnostics Inc.) were tested *in vivo*. Lumason was prepared according to the protocol provided by the manufacturer. NB (200 μL of 4 x 10^11^ NBs/mL) or Lumason (1:4 dilution) was injected via the mouse tail vein (n=3). Nonlinear ultrasound imaging was done on the AplioXG SSA-790A clinical ultrasound imaging system with the same 12 MHz transducers above. The transducer was oriented along the sagittal plane to simultaneously image the kidney and liver of the mouse. Images were acquired in contrast harmonic imaging (CHI) mode with parameters set as: 65 dB dynamic range, 70 dB gain, 0.2 imaging frame rate per second, and peak negative pressure of 245 kPa (equivalent to mechanical index of 0.1). 500 frames (^~^30 mins) of ultrasound images were acquired and the intensity per frame is analyzed with a built-in software (CHI-Q). Enhancement by NBs was calculated by normalizing the measured intensity of the NB solution with respect to the intensity before the injection of NB. *In vivo* half-life and *in vivo* decay onset were calculated by using calculated enhancement and with the formula for a 5-parameter logistic function.

### NB and MicroMarker^®^ stability *in vivo* studies

Mice were handled according to a protocol approved by the Institutional Animal Care and Use Committee (IACUC) at Case Western Reserve University. HCT-15 cells (0.1 mL) in sterile 1X PBS were subcutaneously injected into the dorsal side of BALB/c nude mice at a concentration of 5 x10^6^ cells/mL. Animals were observed every other day until tumors reached at least 8 mm in diameter. NB (200 μL) or Micromarker^®^ (50 μL) was injected via the mouse tail vein. Nonlinear ultrasound imaging was done in VisualSonics Vevo 3100 using MX250 transducer set at 18 MHz, 4% power, 30 dB contrast gain, 28 dB gain, 40 dB dynamic range, 22 frame per second). The entire tumor was then scanned via a 3d motor with respiratory gating at 0.051 mm step size. The 2d slices were reconstructed and analyzed using VevoCQ.

## Supporting Information

Supporting Information is available from the Nanoscale or from the author.

## Acknowledgments

This work was funded by the National Institutes of Health (1R01EB025741-01) and the Office of the Assistant Secretary of Defense for Health Affairs, through the Prostate Cancer Research Program under Award No. W81XWH-16-1-037I and W81XWH-16-1-0372. We also acknowledge additional support from the Case Comprehensive Cancer Center P30CA043703 in the form of a pilot grant and T32 GM008803 training grant support to CCE. Views and opinions of, and endorsements by the author(s) do not reflect those of the National Institutes of Health or of the Department of Defense.

## Notes and References

1 T. Szabo, No TitleDiagnostic Ultrasound Imaging: Inside Out, Academic Press, Second Edi., 2013.

2 R. J. Paproski, A. Forbrich, E. Huynh, J. Chen, J. D. Lewis, G. Zheng and R. J. Zemp, Small, 2016, 12, 371–380.

3 X. Wang, H. Chen, K. Zhang, M. Ma, F. Li, D. Zeng, S. Zheng, Y. Chen, L. Jiang, H. Xu and J. Shi, Small, 2014, 10, 1403–1411.

4 K. P. Hadinger, J. P. Marshalek, P. S. Sheeran, P. A. Dayton and T. O. Matsunaga, Ultrasound Med. Biol., 2018, 44, 2728–2738.

5 J. D. Rojas and P. A. Dayton, Ultrasound Med. Biol., 2019, 45, 177–191.

6 A. G. Nyankima, J. D. Rojas, R. Cianciolo, K. A. Johnson and P. A. Dayton, Ultrasound Med. Biol., 2018, 44, 368–376.

7 Z. Zha, S. Wang, S. Zhang, E. Qu, H. Ke, J. Wang and Z. Dai, Nanoscale, 2013, 5, 3216.

8 F. Dong, J. Zhang, K. Wang, Z. Liu, J. Guo and J. Zhang, Nanoscale, 2019, 11, 1123–1130.

9 Y. Xie, J. Wang, Z. Wang, K. A. Krug and J. D. Rinehart, Nanoscale, 2018, 10, 12813–12819.

10 B. A. Kaufmann, J. R. Lindner, J. C. Wu and J. Narula, Curr. Opin. Biotechnol., 2007, 18, 11–16.

11 M. A. Borden, H. Zhang, R. J. Gillies, P. A. Dayton and K. W. Ferrara, Biomaterials, 2008, 29, 597–606.

12 B. B. Goldberg, J. B. Liu and F. Forsberg, Ultrasound Med. Biol., 1994, 20, 319–333.

13 E. G. Schutt, T. J. Pelura and R. M. Hopkins, Acad. Radiol., 1996, 3 Suppl 2, S188–S190.

14 F. Yang, S. Hu, Y. Zhang, X. Cai, Y. Huang, F. Wang, S. Wen, G. Teng and N. Gu, Adv. Mater., 2012, 24, 5205–5211.

15 P. L. Lin, R. J. Eckersley and E. A. H. Hall, Adv. Mater., 2009, 21, 3949–3952.

16 H. Y. Huang, H. L. Liu, P. H. Hsu, C. S. Chiang, C. H. Tsai, H. S. Chi, S. Y. Chen and Y. Y. Chen, Adv. Mater., 2015, 27, 655–661.

17 Y. Zhou, Z. Wang, Y. Chen, H. Shen, Z. Luo, A. Li, Q. Wang, H. Ran, P. Li, W. Song, Z. Yang, H. Chen, Z. Wang, G. Lu and Y. Zheng, Adv. Mater., 2013, 25, 4123–4130.

18 E. Y. Lukianova-Hleb, X. Ren, J. A. Zasadzinski, X. Wu and D. O. Lapotko, Adv. Mater., 2012, 24, 3831–3837.

19 N. McDannold, N. Vykhodtseva and K. Hynynen, AIP Conf. Proc., 2007, 911, 547–553.

20 E. C. Unger, T. Porter, W. Culp, R. Labell, T. Matsunaga and R. Zutshi, Adv. Drug Deliv. Rev., 2004, 56, 1291–1314.

21 V. Paefgen, D. Doleschel and F. Kiessling, Front. Pharmacol., 2015, 6, 1–16.

22 S. Garg, A. A. Thomas and M. A. Borden, Biomaterials, 2013, 34, 6862–6870.

23 T. Segers, L. De Rond, N. De Jong, M. Borden and M. Versluis, Langmuir, 2016, 32, 3937–3944.

24 A. Katiyar and K. Sarkar, J. Colloid Interface Sci., 2010, 343, 42–47.

25 H. Mulvana, E. Stride, M. X. Tang, J. V. Hajnal and R. J. Eckersley, Ultrasound Med. Biol., 2012, 38, 1097–1100.

26 C. Hernandez, L. Nieves, A. C. De Leon, R. Advincula and A. A. Exner, ACS Appl. Mater. Interfaces, 2018, 10, 9949–9956.

27 R. H. Perera, H. Wu, P. Peiris, C. Hernandez, A. Burke, H. Zhang and A. A. Exner, Nanomedicine Nanotechnology, Biol. Med., 2017, 13, 59–67.

28 V. Uhlendorf, IEEE Trans. Ultrason. Ferroelectr. Freq. Control, 1994, 41, 70–79.

29 K. Ferrara, R. Pollard and M. Borden, Annu. Rev. Biomed. Eng., 2007, 9, 415–447.

30 Y. Murai, Y. Oishi, Y. Takeda and F. Yamamoto, Exp. Fluids, 2006, 41, 343–352.

31 A. Kanai and H. Miyata, Int. J. Numer. Methods Fluids, 2001, 35, 593–615.

32 D. Leguillon and E. Martin, Int. J. Fract., 2013, 179, 157–167.

33 D. Leguillon and E. Martin, Int. J. Fract., 2013, 179, 169–178.

34 B. M. Discher, Y. Won, D. S. Ege, J. C. Lee, F. S. Bates, D. E. Discher and D. A. Hammer, Adv. Sci., 2012, 284, 1143–1146.

35 N. Mohandas and P. G. Gallagher, Blood, 2009, 112, 3939–3948.

36 W. T. S. Huck, Nat. Mater., 2005, 4, 271–272.

37 R. G. S. Maheshwari, R. K. Tekade, P. A. Sharma, G. Darwhekar, A. Tyagi, R. P. Patel and D. K. Jain, Saudi Pharm. J., 2012, 20, 161–170.

38 G. M. El Maghraby, B. W. Barry and A. C. Williams, Eur. J. Pharm. Sci., 2008, 34, 203–222.

39 W. Terakosolphan, J. L. Trick, P. G. Royall, S. E. Rogers, O. Lamberti, C. D. Lorenz, B. Forbes and R. D. Harvey, Langmuir, 2018, 34, 6941–6954.

40 A. C. Biondi and E. A. Disalvo, Biochim. Biophys. Acta - Biomembr., 1990, 1028, 43–48.

41 L. Pocivavsek, K. Gavrilov, K. D. Cao, E. Y. Chi, D. Li, B. Lin, M. Meron, J. Majewski and K. Y. C. Lee, Biophys. J., 2011, 101, 118–127.

42 C. Hernandez, S. Gulati, G. Fioravanti, P. L. Stewart and A. A. Exner, Sci. Rep., 2017, 7, 13517.

43 J. Lee, W. Shen, K. Payer, T. P. Burg and S. R. Manalis, Nano Lett., 2010, 10, 2537–2542.

44 M. Godin, A. K. Bryan, T. P. Burg, K. Babcock and S. R. Manalis, Appl. Phys. Lett., 2007, 91, 1–4.

45 T. P. Burg, M. Godin, S. M. Knudsen, W. Shen, G. Carlson, J. S. Foster, K. Babcock and S. R. Manalis, Nature, 2007, 446, 1066–1069.

46 S. Olcum, N. Cermak, S. C. Wasserman, K. S. Christine, H. Atsumi, K. R. Payer, W. Shen, J. Lee, A. M. Belcher, S. N. Bhatia and S. R. Manalis, Proc. Natl. Acad. Sci., 2014, 111, 1310–1315.

47 C. Hernandez, E. C. Abenojar, J. Hadley, A. C. de Leon, R. Coyne, R. Perera, R. Gopalakrishnan, J. P. Basilion, M. C. Kolios and A. A. Exner, Nanoscale, 2019, 11, 851–855.

48 T. Segers, D. Lohse, M. Versluis and P. Frinking, Langmuir, 2017, 33, 10329–10339.

49 S. Duangjit, Y. Obata, H. Sano, Y. Onuki, P. Opanasopit, T. Ngawhirunpat, T. Miyoshi, S. Kato and K. Takayama, Biol. Pharm. Bull., 2014, 37, 239–247.

50 J. Majewski, T. L. Kuhl, K. Kjaer and G. S. Smith, Biophys. J., 2001, 81, 2707–2715.

51 E. H. Lee, A. Kim, Y. K. Oh and C. K. Kim, Biomaterials, 2005, 26, 205–210.

52 R. Alvarez-Román, A. Naik, Y. N. Kalia, R. H. Guy and H. Fessi, Pharm. Res., 2004, 21, 1818–1825.

53 M. Trotta, E. Peira, M. E. Carlotti and M. Gallarate, Int. J. Pharm., 2004, 270, 119–125.

54 R. M. Elmoslemany, O. Y. Abdallah, L. K. El-Khordagui and N. M. Khalafallah, AAPS PharmSciTech, 2012, 13, 723–731.

55 K. Sato and S. Ueno, Curr. Opin. Colloid Interface Sci., 2011, 16, 384–390.

56 S. Hoche, M. A. Hussein and T. Becker, Ultrasonics, 2015, 57, 65–71.

57 K. Zell, J. I. Sperl, M. W. Vogel, R. Niessner and C. Haisch, Phys. Med. Biol., DOI:10.1088/0031-9155/52/20/N02.

58 H. W. Persson and C. H. Hertz, Ultrasonics, 1985, 23, 83–89.

59 O. Falou, A. Jafari Sojahrood, J. Carl Kumaradas and M. C. Kolios, J. Acoust. Soc. Am., 2012, 132, 1820–1829.

60 O. Falou, C. Church, R. E. Baddour, G. Nathanael, C. Church, G. J. Czarnota, C. Church, J. C. Kumaradas, M. C. Kolios and C. Church, J. Acoust. Soc. Am., 2008, 124, EL278–EL283.

61 R. E. Baddour, M. D. Sherar, J. W. Hunt, G. J. Czarnota and M. C. Kolios, J. Acoust. Soc. Am., 2005, 117, 934–943.

62 T. van Rooij, Y. Luan, G. Renaud, A. F. W. van der Steen, M. Versluis, N. de Jong and K. Kooiman, Ultrasound Med. Biol., 2015, 41, 1432–1445.

63 P. Marmottant, S. van der Meer, M. Emmer, M. Versluis, N. de Jong, S. Hilgenfeldt and D. Lohse, J. Acoust. Soc. Am., 2005, 118, 3499–3505.

64 J. S. Lum, J. D. Dove, T. W. Murray and M. A. Borden, Langmuir, 2016, 32, 9410–9417.

65 S. Alonso-Romanowski, A. C. Biondi and E. A. Disalvo, J. Membr. Biol., 1989, 108, 1–11.

66 J. H. Crowe, L. M. Crowe and D. Chapman, Science (80-.)., 1984, 223, 701–703.

67 J. M. Boggs and G. Rangaraj, Biochim. Biophys. Acta - Biomembr., 1985, 816, 221–233.

68 R. H. Abou-Saleh, J. R. McLaughlan, R. J. Bushby, B. R. Johnson, S. Freear, S. D. Evans and N. H. Thomson, Langmuir, 2019, acs.langmuir.8b04130.

69 E. Huynh, B. Y. C. Leung, B. L. Helfield, M. Shakiba, J. A. Gandier, C. S. Jin, E. R. Master, B. C. Wilson, D. E. Goertz and G. Zheng, Nat. Nanotechnol., 2015, 10, 325–332.

70 V. Sboros, Adv. Drug Deliv. Rev., 2008, 60, 1117–1136.

71 I. López-Montero, L. R. Arriaga, F. Monroy, G. Rivas, P. Tarazona and M. Vélez, Langmuir, 2008, 24, 4065–4076.

72 U. G. K. Wegst, H. Bai, E. Saiz, A. P. Tomsia and R. O. Ritchie, Nat. Mater., 2015, 14, 23–36.

73 T. Yin, P. Wang, R. Zheng, B. Zheng, D. Cheng, X. Zhang and X. Shuai, Int. J. Nanomedicine, 2012, 7, 895–904.

74 Y. Gao, C. Hernandez, H. X. Yuan, J. Lilly, P. Kota, H. Zhou, H. Wu and A. A. Exner, Nanomedicine Nanotechnology, Biol. Med., 2017, 13, 2159–2168.

75 K. Maruyama, S. Okuizumi, O. Ishida, H. Yamauchi, H. Kikuchi and M. Iwatsuru, Int. J. Pharm., 1994, 111, 103–107.

76 C. C. Chen, S. R. Sirsi, S. Homma and M. A. Borden, Ultrasound Med. Biol., 2012, 38, 492–503.

77 Z. Zhang, J. Xu, B. Liu, F. Chen, J. Li, Y. Liu, J. Zhu and C. Shen, Inflammopharmacology, DOI:10.1007/s10787-018-0534-5.

78 A. Sartore-Bianchi, S. Marsoni and S. Siena, JAMA Oncol., 2017, 4, 2017–2018.

79 Y. Nasseri and S. J. Langenfeld, Surg. Clin. North Am., 2017, 97, 503–513.

80 E.-T. Cartana, D. Gheonea, I. Cherciu, I. Streaţa, C.-D. Uscatu, E.-R. Nicoli, M. Ioana, D. Pirici, C.-V. Georgescu, D.-O. Alexandru, V. Şurlin, G. Gruionu and A. Săftoiu, Endosc. Ultrasound, 2018, 7, 175.

