## Supporting Information for "Bubble Trouble: Conquering Microbubble Limitations in Contrast Enhanced Ultrasound Imaging by Nature-Inspired Ultrastable Echogenic Nanobubbles"

| Reagents (for 5 mL solution) | PG-PL | PG-Gly-PL | Gly-PL | PL |
|---|---|---|---|---|
| C22 (DBPC) | 30.5 mg | 30.5 mg | 30.5 mg | 30.5 mg |
| DPPA | 5.0 mg | 5.0 mg | 5.0 mg | 5.0 mg |
| DPPE | 10.0 mg | 10.0 mg | 10.0 mg | 10.0 mg |
| DSPE-mPEG 2000 | 5.0 mg | 5.0 mg | 5.0 mg | 5.0 mg |
| PBS | 4.0 mL | 4.0 mL | 4.0 mL | 5.0 mL |
| **Glycerol** | **0** | **0.5 mL** | **1.0 mL** | **0 mL** |
| **Propylene Glycol** | **1.0 mL** | **0.5 mL** | **0 mL** | **0 mL** |

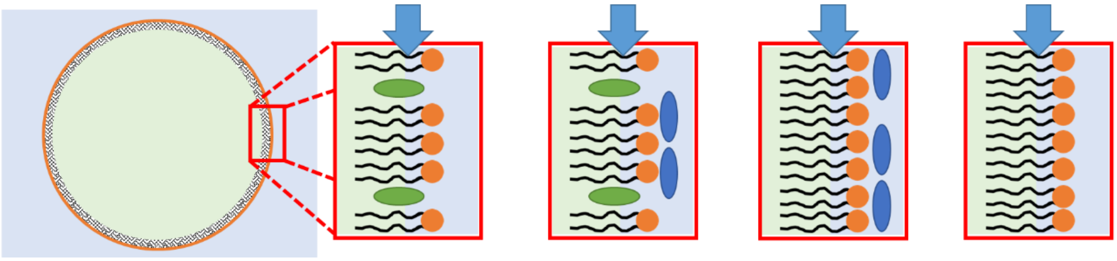

**Table S1.** Chemical composition of NBs and schematic representation of NB shell composition

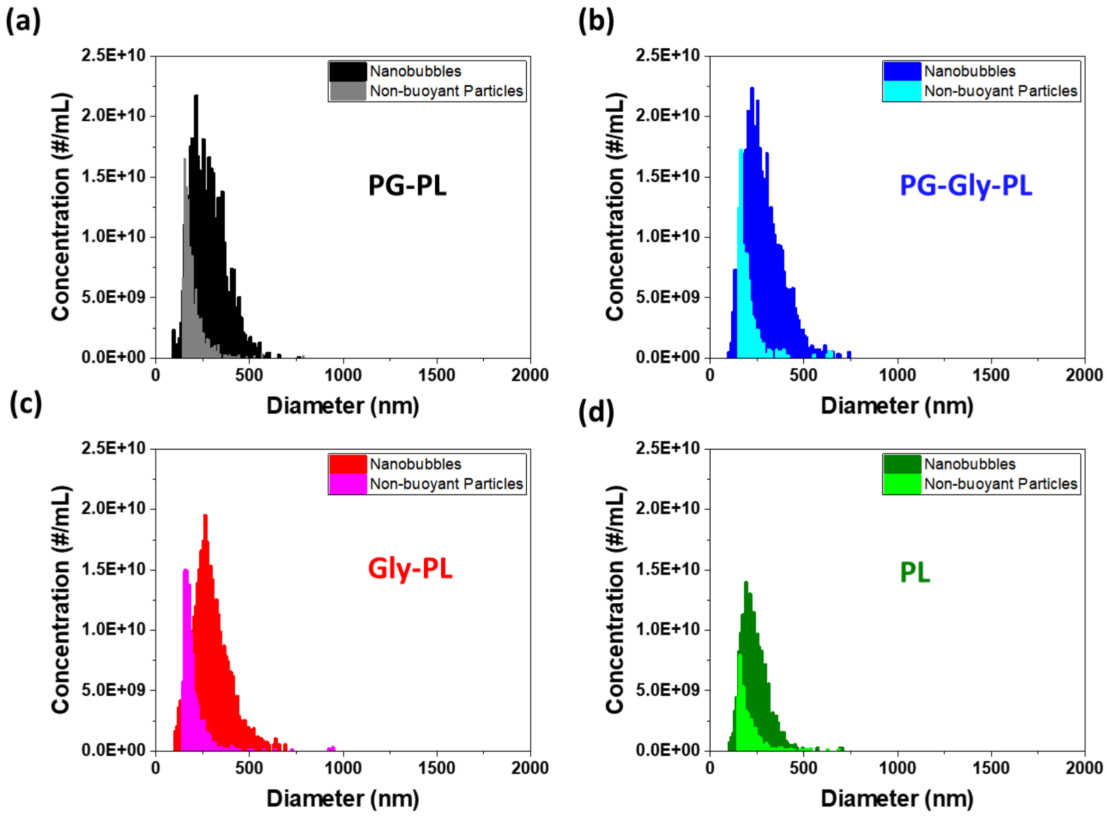


**Figure S1.** Particle size distribution of NBs and non-buoyant particles measured by resonant mass measurement.

| Sample | Zeta Potential (mV) |
|---|---|
| PG-PL | -2.53 ± 0.88 |
| PG-Gly-PL | -2.15 ± 1.78 |
| Gly-PL | -2.04 ± 1.70 |

**Table S2.** Zeta potential of different NB solutions. Mean ± SE (n=3).

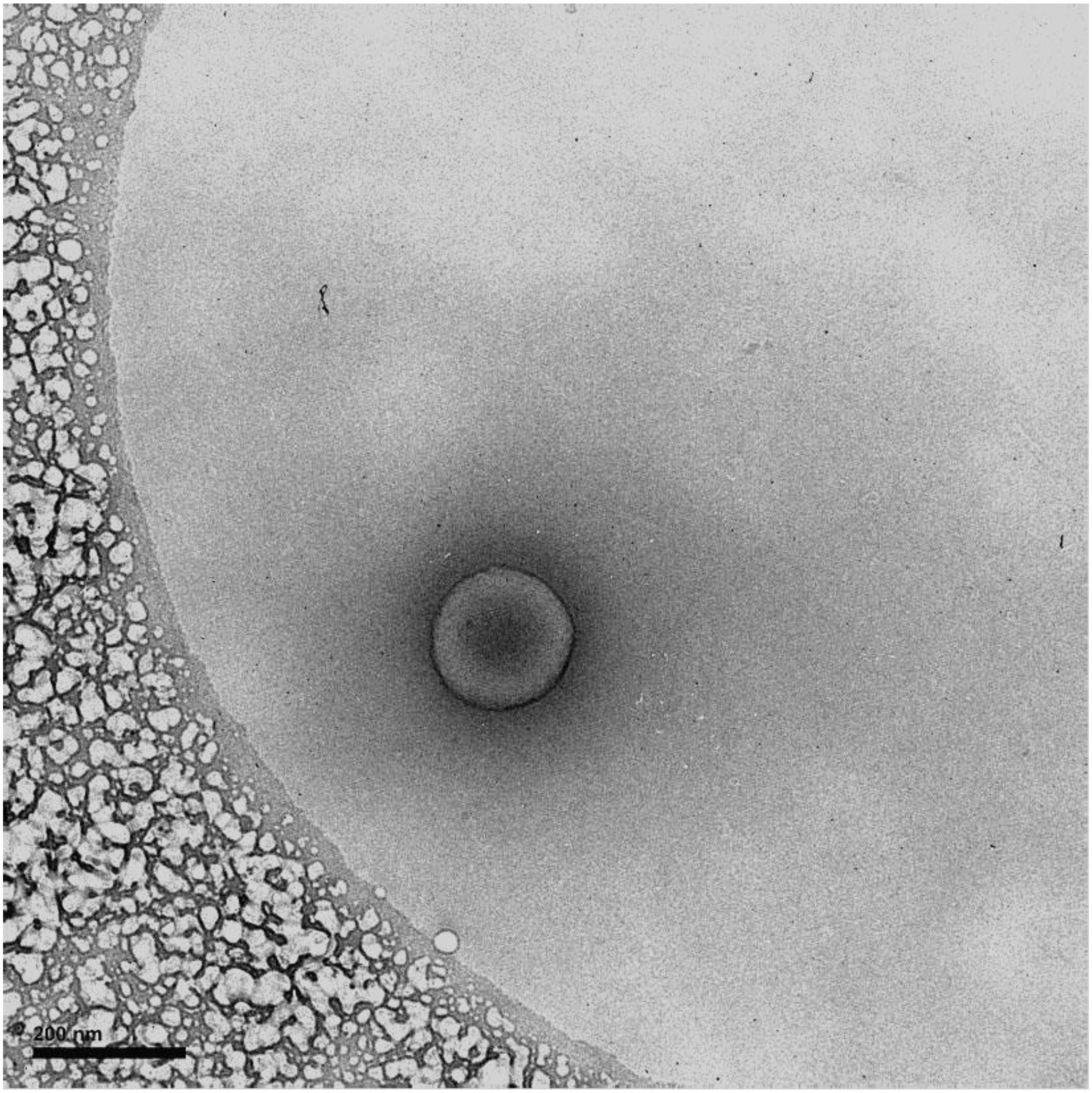


Figure S2. Cryo-electron microscopy (Cryo-EM) image of PG-Gly-PL showing the nanobubble membrane and the dense $C_3F_8$ gas core.

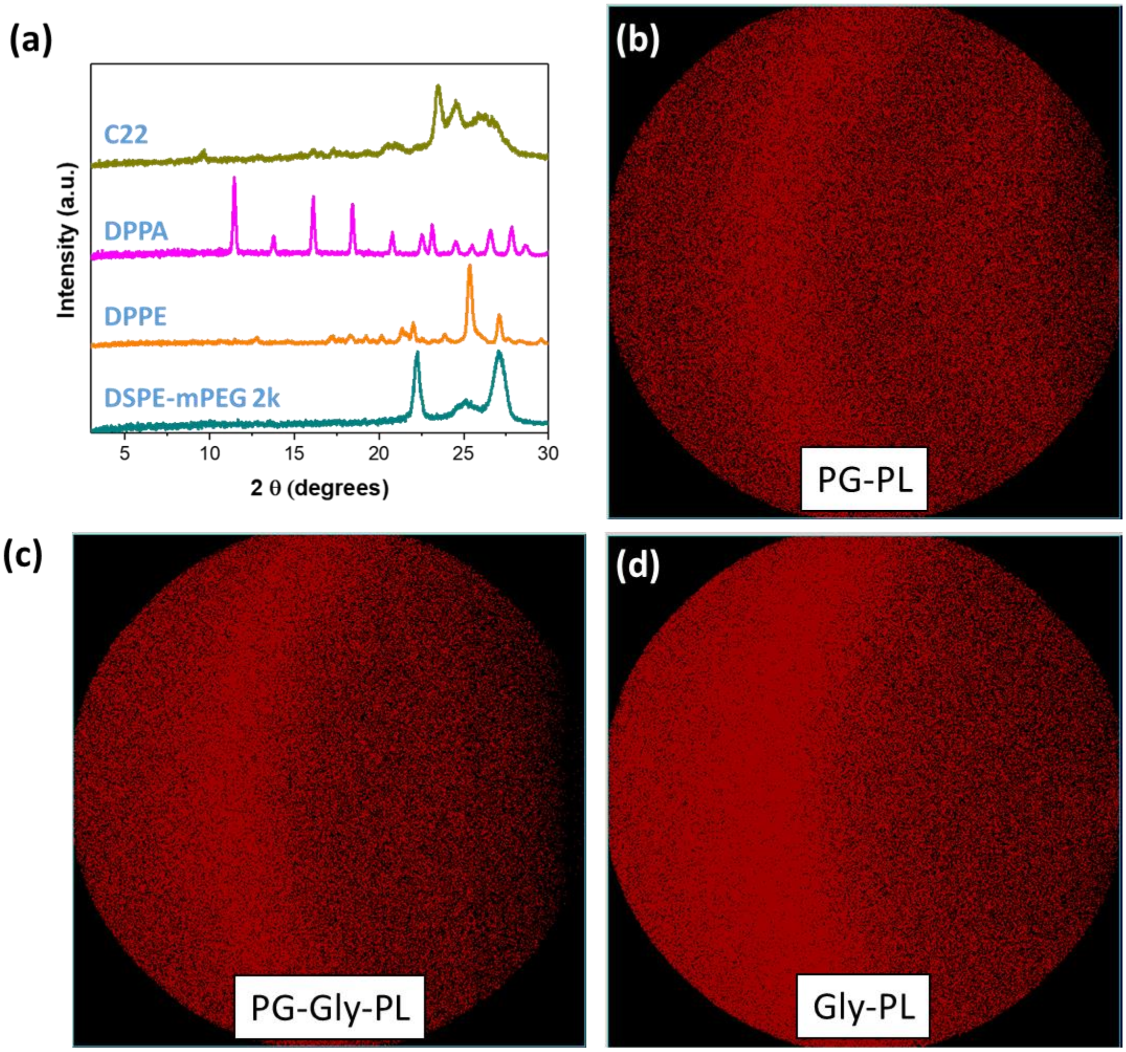


**Figure S3.** X-ray diffractograms of pure phospholipids used in the NB formulation and 2D XRD of freeze-dried NB membrane.

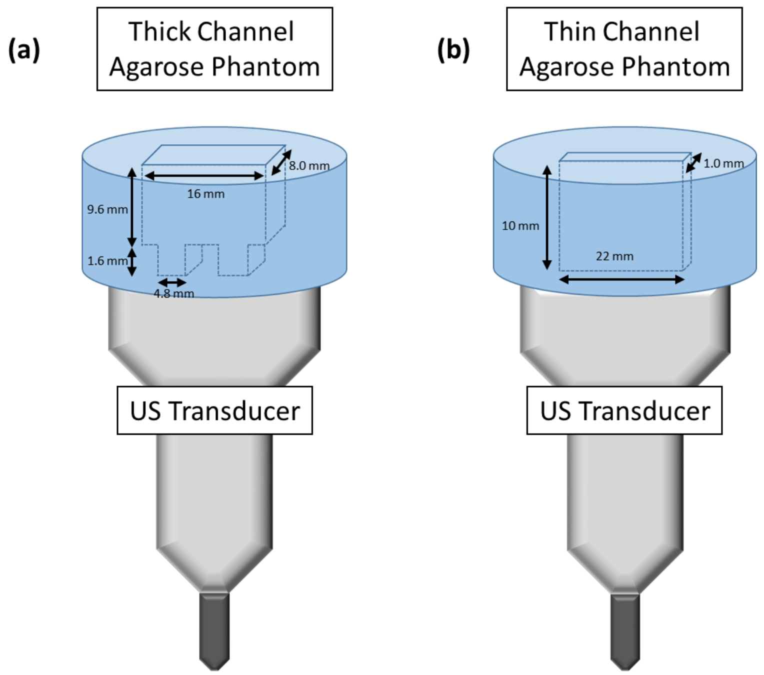


**Figure S4.** Schematic diagram showing the dimensions and orientation of the tissue-mimicking agarose phantoms used for echogenicity (a) and stability experiments (b).

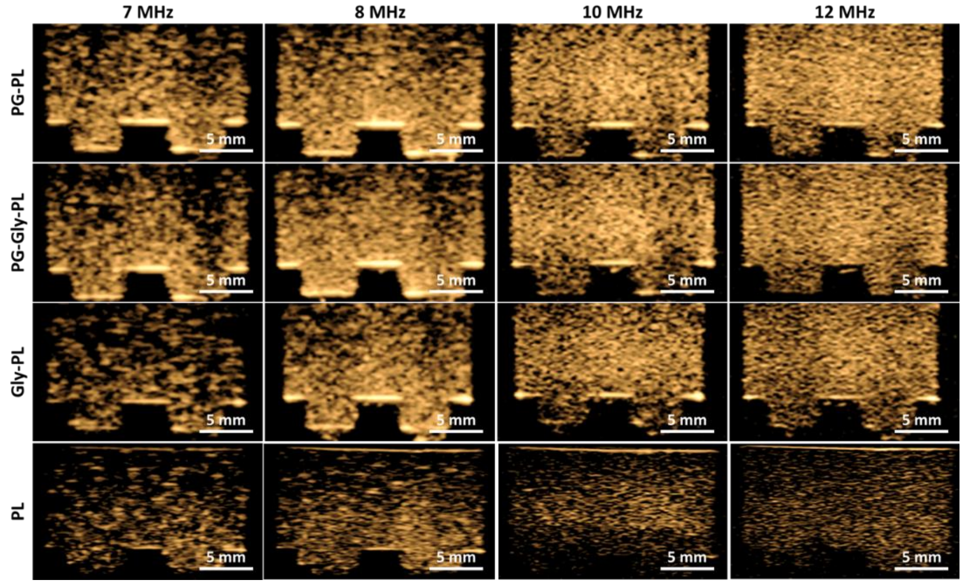


**Figure S5.** Ultrasound images captured in contrast harmonic imaging mode of NB solutions acquired at different frequencies.

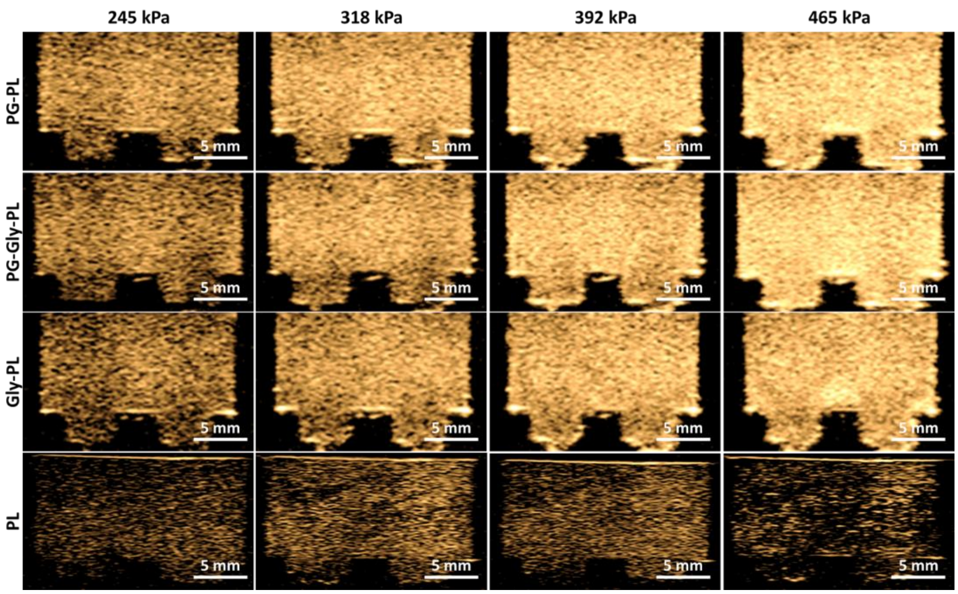


**Figure S6.** Ultrasound images captured in contrast harmonic imaging mode of NB solutions acquired at different peak negative pressures.

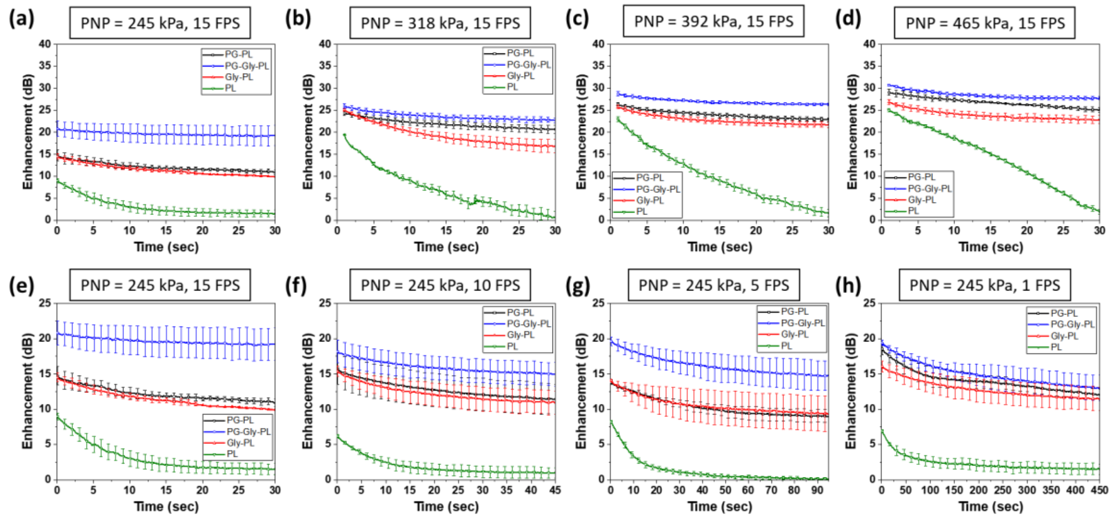


**Figure S7.** Stability curves of NB solutions acquired at different peak negative pressure and US pulse rate. Mean ± SE (n=3).

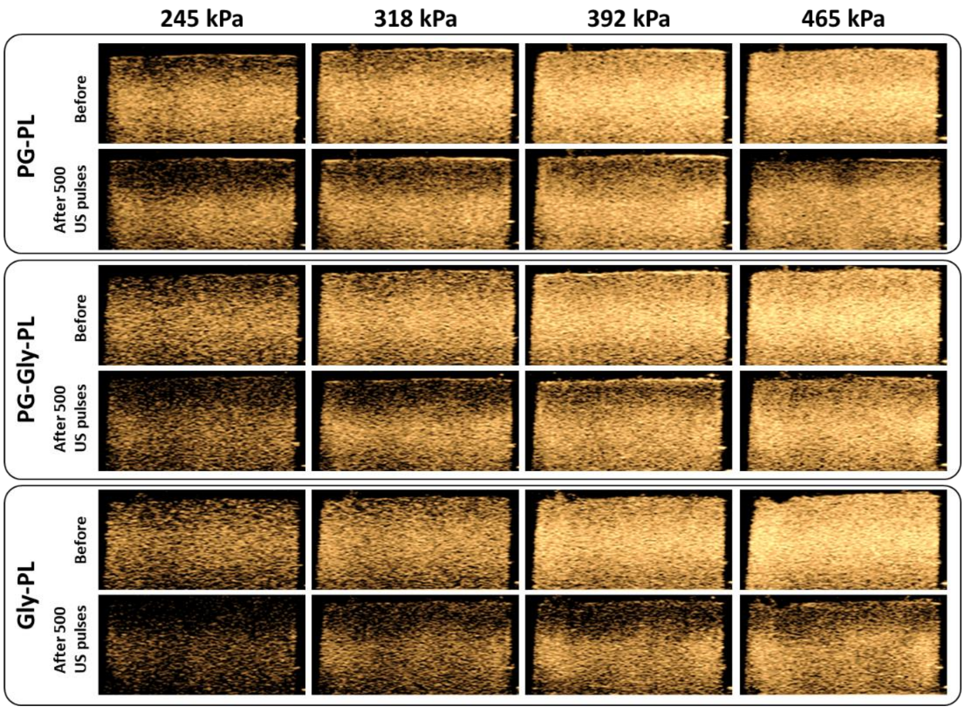


**Figure S8.** Ultrasound images captured in contrast harmonic imaging mode before and after exposure to 500 frames of varying peak negative pressures.

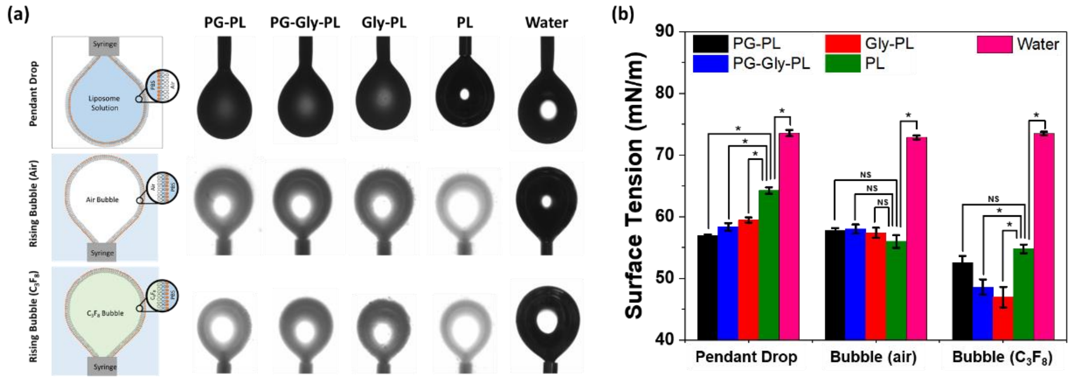


**Figure S9.** Surface tension measurement of NB solutions acquired via the pendant drop method, rising bubble method in air, and rising bubble method in $C_3F_8$. Mean ± SE (n = 3).

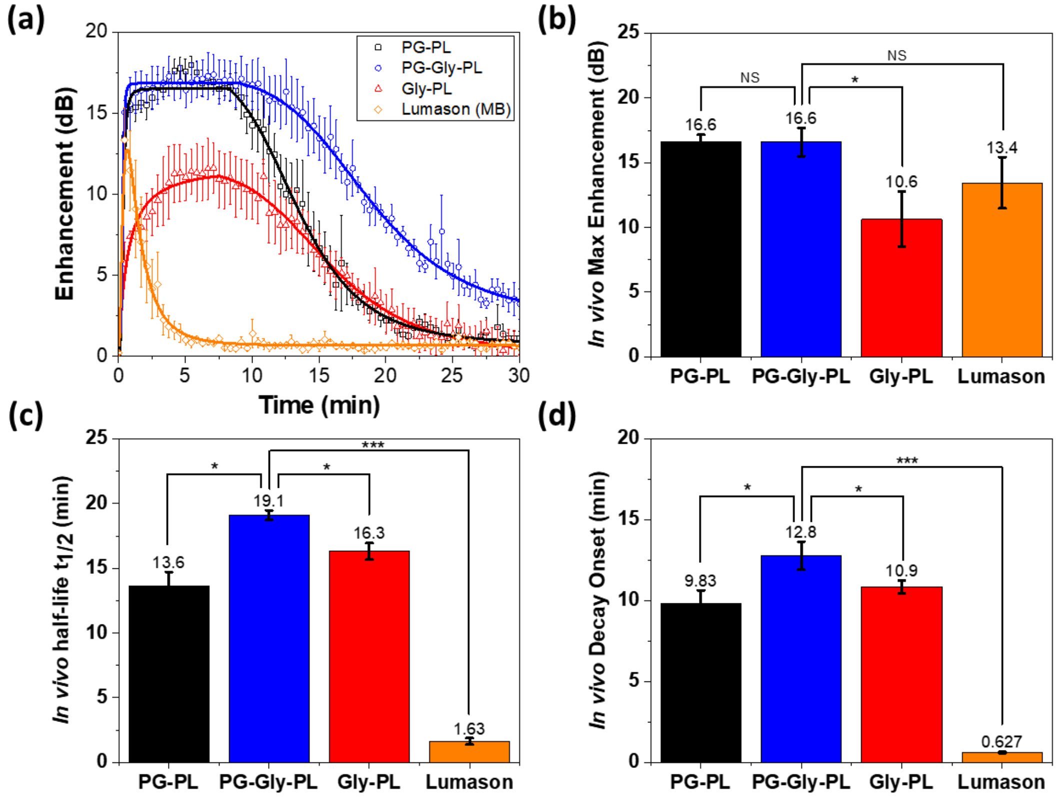


**Figure S10.** *In vivo* stability of NBs and Lumason in mouse liver showing similar behavior to that in the mouse kidney despite the presence of liver macrophages. Mean ± SE (n = 3)

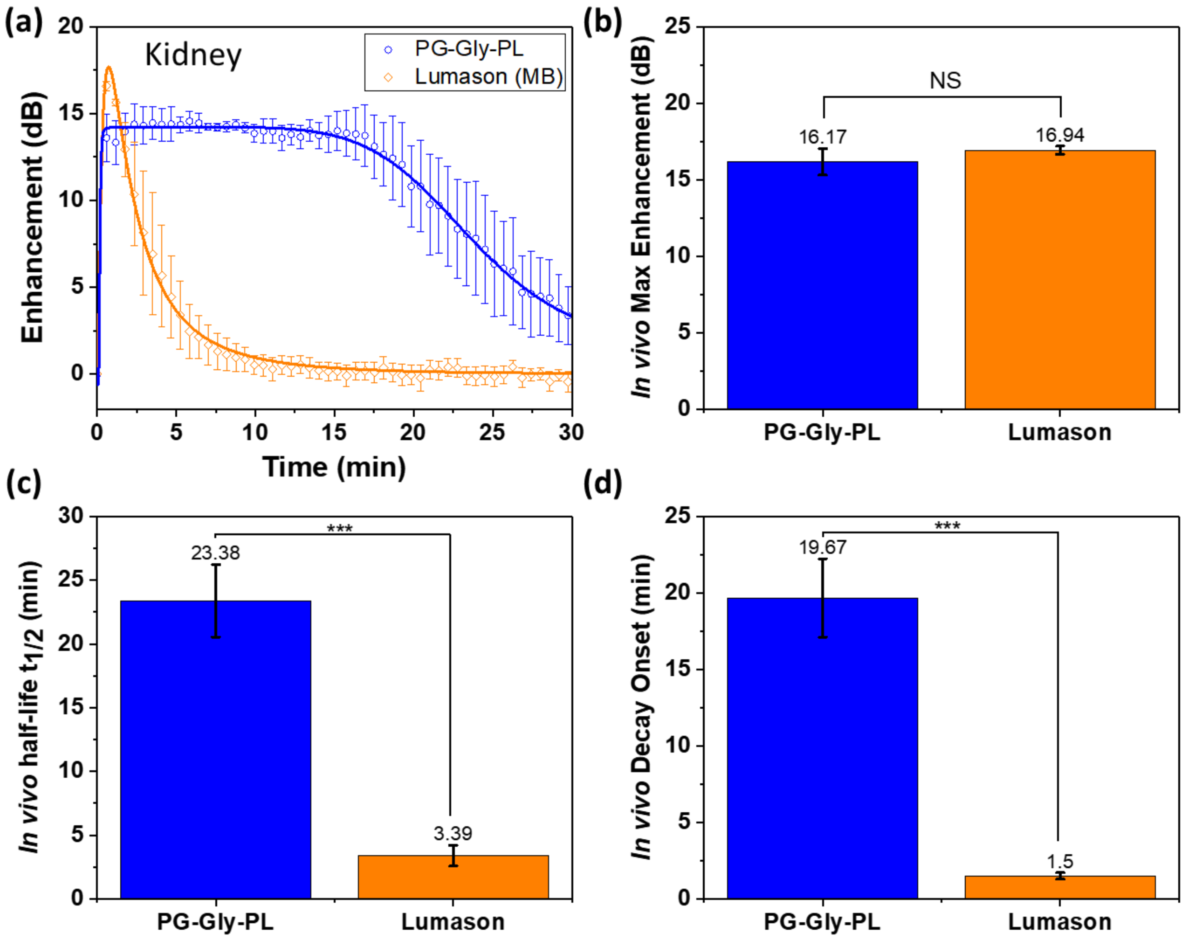


**Figure S11.** *In vivo* stability of PG-Gly-PL and Lumason in the kidney of a mouse with colorectal flank tumor. Mean ± SE (n = 3)

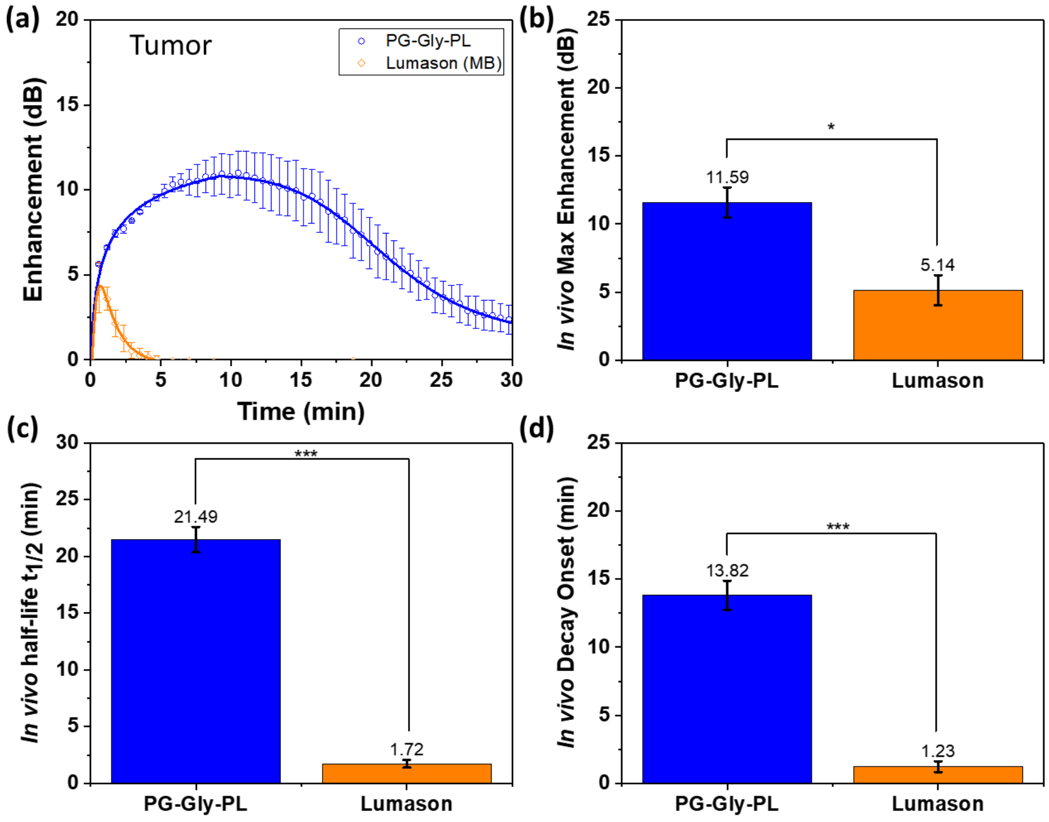


**Figure S12.** *In vivo* stability of PG-Gly-PL and Lumason in the colorectal flank tumor of a mouse. Mean ± SE (n = 3)
